# A semiparametric method to test for correlated evolution in a phylogenetic context

**DOI:** 10.1101/2025.02.20.639307

**Authors:** Luke J. Harmon, Liam J. Revell

## Abstract

Phylogenetic comparative methods are a broad suite of approaches for studying trait and species diversification using evolutionary trees. In spite of their substantial growth in sophistication and complexity in recent years, among the most commonly-employed phylogenetic comparative method continues to be a simple measure of the evolutionary correlation between variables, while accounting for the statistical non-independence in our data that arises from common descent. The standard parametric phylogenetic approach for measuring the evolutionary correlation between continuously valued characters assumes a model called Brownian motion for the evolution of our traits. Here, we introduce a new semiparametric method that relaxes this assumption by testing for the evolutionary correlation between variables based on contrast ranks, and then obtains a null distribution on the test statistic via random permutation. We show that this approach has reasonable statistical properties: type I error close to the nominal level, and power that is similar to fully parametric methods. We conclude by comparing our new method to related approaches.

## 1 INTRODUCTION

Statistical phylogenetic comparative methods have become an invaluable tool in contemporary evolutionary research (Felsenstein 1985; Revell and Harmon 2022). This group of methodologies comprises a large and growing set of approaches for studying increasingly sophisticated evolutionary hypotheses of trait and species diversification using phylogenetic trees (reviewed in Harvey and Pagel 1991; Nunn 2011; O’Meara 2012; Harmon 2019; Revell and Harmon 2022). Nonetheless, among the most common evolutionary hypothesis studied by researchers using statistical comparative methods is the simple question of whether one trait is correlated with another – while accounting for the non-independence of the species that make up our data points due to their shared histories (Felsenstein 1985; reviewed in Nunn 2011; Harmon 2019; Revell and Harmon 2022).

When measuring and undertaking a hypothesis test of this nature, phylogenetic comparative methods typically assume a particular underlying model of trait evolution, such as the Brownian motion model (Felsenstein 1973; Felsenstein 1985; O’Meara et al. 2006). Brownian motion (also called a Wiener process) is a continuous-time stochastic diffusion process in which the expected value is constant and the variance increases as a linear function of time multiplied by the Brownian motion rate, typically denoted *σ* 2 (Felsenstein 1973; Felsenstein 2004; O’Meara et al. 2006; Harmon 2019; Revell and Harmon 2022). In particular, under the phylogenetically independent contrasts algorithm of Felsenstein (1985), both the internal node values in the contrasts algorithm, and the variances of the (non-standardized) contrasts, are calculated assuming a Brownian evolutionary process on the tree (Felsenstein 1985, 2004; Harmon 2019). Likewise, under phylogenetic generalized least squares regression (PGLS, Grafen 1989; Rohlf 2001), we assume that either the data themselves, or their residual errors, are autocorrelated in a manner consistent with Brownian motion evolution– or a closely-related process (e.g., Pagel 1999; Garland and Ives 2000; Rohlf 2001; Revell 2010). Although starting from different assumptions, PICs and PGLS are equivalent (Blomberg et al. 2012). Though there are a number of circumstances in which continuously valued phenotypic traits might genuinely evolve via a Brownian process (e.g., Lynch 1990), violations of the Brownian model are likely be common in real data sets (Blomberg et al. 2003; Freckleton and Harvey 2006; Hunt 2006; Estes and Arnold 2007; Uyeda et al. 2011). Brownian motion has nonetheless persisted as one of the most commonly used trait evolution models for studying evolutionary correlations, largely due to its simplicity, tractability, and apparent robustness to deviations from model assumptions under some circumstances (e.g., Garland et al. 1993; Rohlf 2001; Stone 2011).

Nonetheless, a number of different approaches have been devised to identify violations of Brownian motion in our data while measuring evolutionary correlations, and then to account for these deviations statistically.

First, we can apply various diagnostic tests for independent contrasts that are sensitive to violations of the Brownian model (e.g., Garland 1992; Blomberg et al. 2003; Freckleton and Harvey 2006). For example, if the generative model of the data is Brownian motion, then standardized independent contrasts of Felsenstein (1985) should be independently and identically distributed (i.i.d.). If, on the other hand, standardized contrasts are correlated with their expected variances (that is, the amount of evolutionary branch length that subtends each phylogenetic contrast), this suggests that there may be either more or less evolutionary change along long branches of the phylogeny (depending on the sign of the correlation) than would have been expected under a Brownian process (Garland 1992). We might then attempt various transformations of the branch lengths of the tree so that our contrasts no longer violate this test (Garland 1992).

Another similar and closely-related diagnostic is the node-height test (Freckleton and Harvey 2006). Here, we measure for a relationship between our standardized contrasts and the height (above the root) of the node over which each contrast was calculated (Freckleton and Harvey 2006). Once again, this correlation should be statistically indistinguishable from zero if the Brownian motion model assumption holds for our data.

Both the measures of Garland (1992) and Freckleton and Harvey (2006)’s node-heights test can also be interpreted as measurements of model adequacy. If they fail (that is, if we reject a null hypothesis of no correlation between the standardized contrast and its variance or node height), then this suggests that our underlying assumption of Brownian motion may be invalid (Pennell et al. 2015).

Under circumstances in which Brownian motion has been rejected via a diagnostic test, we might then proceed to consider one or more of a relatively limited set of alternative trait evolution models (e.g., Pagel 1999; Revell 2010). In particular, we may proceed to fit our correlation assuming a different evolutionary model, such as the Ornstein-Uhlenbeck or a speciational trait evolution models (e.g., Hansen 1997; Martins and Hansen 1997). Alternatively, we could transform the branches of the phylogenetic tree, or the elements of the variance-covariance matrix derived from the tree (Grafen 1989; Garland 1992; Pagel 1999; Revell 2010). Nevertheless, there is no general solution to the problem of evolutionary model assumption violations, and on occasion there may simply be no way to coax our data set or tree to fit the assumptions of our analysis!

In statistics, nonparametric tests can sometimes provide robustness in cases where models are not known or provide poor fits to data (e.g., Wasserman 2006). As comparative methods are employed to tackle a wider and wider range of problems, from genomics to cancer biology to evolutionary studies at the broadest scale, there is a growing need for precisely this kind of robust approach. Among existing comparative methods, nonparametric approaches are already available for the analysis of discrete traits (e.g., Maddison 1990), and for morphometric shape data (Adams 2014). To date, however, the only non- or semiparametric comparative method designed specifically for measuring the correlation of continuously valued characters is Ackerly’s (2000) contrast sign test (CST). This test involves comparing the signs (positive or negative) of phylogenetically independent contrasts for two characters (Ackerly 2000). If the two characters are evolving independently, then the number of pairs of contrasts with the same sign should follow a binomial distribution with probability of success *p* = 0.5. The CST provides a robust alternative to fully parametric comparative methods, but is not commonly used. Its primary drawback is that the test can have very low power– in part because it relies only on the contrast signs, and discards all information about their relative magnitudes.

Here we describe a new semiparametric comparative method, that we call the phylogenetic rank correlation (PRC) test. This test, which is based on the correlation of ranks of independent contrasts on a phylogenetic tree, seems to be robust to substantial deviations from a strict assumption of Brownian motion evolution of our traits, but nonetheless has power comparable to standard independent contrasts and much greater than the CST. Consequently, we argue that the PRC test may provide a robust alternative to fully parametric methods, such as PGLS and independent contrasts, when the Brownian motion evolution model is clearly violated for our data (or their residual errors, Rohlf 2001; Revell 2010).

## 2 DESCRIPTION OF THE METHOD

The PRC test has a total of six steps (Figure 1), as follows.

**Figure 1.**
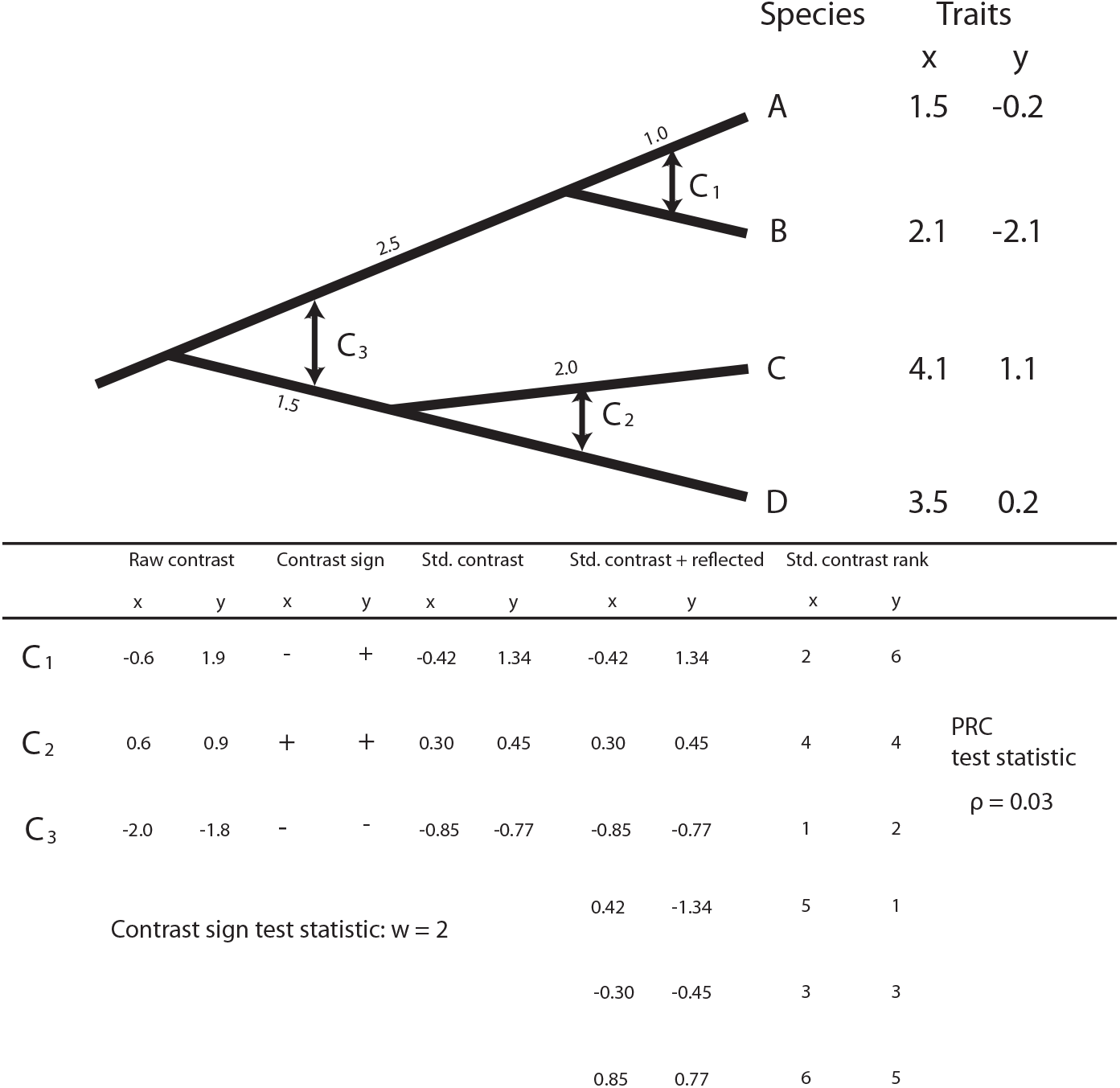
Illustration of the method of phylogenetic rank contrasts, PRC. First, we compute raw independent contrasts in the standard way. For a bifurcating tree with *N* tips there should be *m* = *N* − 1 such contrasts, in which *m* is also the number of nodes in the tree. Next, we standardize the contrasts by their expected variance under Brownian motion following Felsenstein (1985). We duplicate all pairs of contrasts for each of our two variable, but swap the signs (i.e., rotating around the origin, [0, 0]). Next, we convert all contrasts (and their reflections) to ranks. Finally, we calculate the test-statistic, in this case, Spearman’s *ρ*. A P-value for *ρ* is obtained via permutation of the contrasts for either *X* or *Y* (it doesn’t matter which). See main text for more details.

1. Calculate standardized independent contrasts (Felsenstein 1985) for two continuously distributed traits, *X* and *Y* . These contrasts each consist of two numeric values (a contrast for *X* and a contrast for *Y*), and they should number **C**_1_, **C**_2_, through **C**_*m*_ for *m* internal nodes in the tree (Figure 1).
2. Reflect the contrasts, multiplying both the *X* and *Y* value of each pair of contrasts in the two traits by −1. The data will now include two ‘copies’ of each *X*, *Y* pair of contrasts, one with the signs of each contrast reversed. Thus, for example, if contrast **C**_1_ consists of the numeric values [−0.6, 1.9], we would create an additional contrast in our data set, **C**_*m*+1_, that was equal to [0.6, − 1.9] (Figure, and then proceed to do the same for all *m* contrasts in the tree. This reflection is necessary due to the arbitrary direction of subtraction chosen when calculating contrasts, as discussed below.
3. Create ranks independently for the numeric values of the contrasts for each variable, *X* and *Y* (Figure 1). In the case of tied values, we should assign the mean rank to all contrast values in the tie.
4. Calculate the rank correlation test statistic. Here, we use Spearman’s rank correlation coefficient (Spearman’s *ρ*, Spearman 1904), but we could have also use Kendall’s *τ* (Kendall 1938).
5. Generate a null distribution for the test statistic via randomization of the contrasts. To do this, we must permute the numeric values of the undoubled independent contrasts for one variable, *X* or *Y* (it doesn’t matter which). Then we reflect the contrasts, create new ranks, and calculate the test statistic. (Note that the ranks of the variable that is not permuted will not change!) We repeat this permutation procedure a large number of times (say, 999 or 9,999), each time recalculating the test statistic (*ρ* or *τ*) to obtain a null distribution.
6. Compare the test statistic with the null distribution from permutation to obtain a P-value of the test. We calculate our P-value as 2× the fraction of permuted test-statistic values (for a two-tailed test) that are equally or more extreme than our measured value of *ρ* (or *τ*).

Several of these steps might require some additional comment or explanation. For step (1.), standardized contrasts are calculated. The calculation of these contrasts, as originally described by Felsenstein (1985), assumes a Brownian motion model of evolution; however, since the standardized contrasts are to be converted into ranks, we maintain that violating this assumption will have a much smaller effect on their subsequent analysis than in a fully parametric approach (see below). In practice, by converting contrasts into ranks we have assumed only that contrasts subtending longer branches of the tree tend to be larger in absolute magnitude. This assumption is compatible with a broad range of evolutionary scenarios, as long as variance among species tends to increase through time (see Estes and Arnold 2007).

In step (2.), reflecting the contrasts is critical because the direction of any particular contrast is arbitrary. (This can be understood via the thought experiment in which we rotate descendant edges around any node of the tree. Positively-valued contrasts in *X* or *Y* will become negative, as well as the converse.) Reflecting contrasts around the origin ensures that our analysis is unchanged by node rotations of the tree.

This procedure is likewise conceptually equivalent to forcing our regression line through the point [0, 0], which is required when testing for correlations of independent contrasts (Garland et al. 1992). We do not have to worry about inflating degrees of freedom by doubling the number of data points because statistical significance is to be determined by permutations.

Finally, in step (4.), any nonparametric test statistic could be used for this test (see, e.g., Wasserman 2006). For simplicity, we have chosen to use Spearman’s rank correlation (Spearman’s *ρ*), which is simply the Pearson product-moment correlation of the ranks of the two variables (Spearman 1904). Kendall’s *τ* (Kendall 1938) is another viable option that could be substituted with no other changes to the described methodology.

We provide a worked example of the calculations involved in this test in Figure 1. Software to carry out the PRC test, written in the R statistical computing language (R Core Team 2024), will be made publicly available as part of the popular *geiger* R phylogenetics package (Harmon et al. 2008; Pennell et al. 2014).

## 3 STATISTICAL PROPERTIES OF THE METHOD

To explore the statistical properties of our semiparametric evolutionary correlation test, the PRC, we measured type I error and power of the test under a range of circumstances. Then we compared both to the same measures for both non-phylogenetic (i.e., OLS or ‘standard’ non-phylogenetic correlation) and related phylogenetic comparative methods (independent contrasts, contrasts with a statistical distribution obtained via permutation, and the contrasts sign test of Ackerly 2000).

To measure type I error we simulated data for *X* and *Y* under Brownian motion evolution and with no evolutionary correlation between the two traits. For this simulation, we first began by generating 1,000 pure-birth (i.e., Yule) phylogenetic trees containing from *N* = 10 to 100 taxa on ten taxon intervals (i.e., *N* = 10, 20, 30, etc.), an exercise that resulted in a total of 9 × 1, 000 = 9, 000 simulated trees. These trees were rescaled to have a total height of 1.0, and each of *X* and *Y* were simulated (independently, as they are uncorrelated) with a Brownian motion rate of *σ* 2 = 1.0.

Using this set of 9,000 trees and simulated datasets for *X* and *Y*, we then proceeded to fit a standard (OLS) regression (“Standard”), a contrasts regression (“IC”), a contrasts regression but in which a P-value for hypothesis testing was obtained via random permutation (“IC Perm”), the contrasts sign test (“CST”), and our phylogenetic rank correlation (“PRC”). We decided to include a permutation test based on phylogenetically independent contrasts to ensure that our measured type I error and power for the PRC were due to the use of a rank correlation, rather than for having substituted a permutation distribution for the parametric distribution typically used in phylogenetic correlation with independent contrasts. For each analysis we computed a P-value, and then we estimated the type I error rate as the fraction of P-values from each set of simulation conditions and analysis with values equal to or smaller than 5%.

Results from this analysis are given in Figure 2a. Consistent with expectations (e.g., Revell 2010), we found high type I error of standard, non-phylogenetic correlation for phylogenetic data simulated under Brownian motion (Figure 2a), at a rate that increased with increasing numbers of taxa, *N* (Figure 2a). By contrast, when the assumption of Brownian motion evolution held true, all phylogenetic analyses had type I error rates close to the nominal level of *α* = 0.05 (Figure 2a).

**Figure 2.**
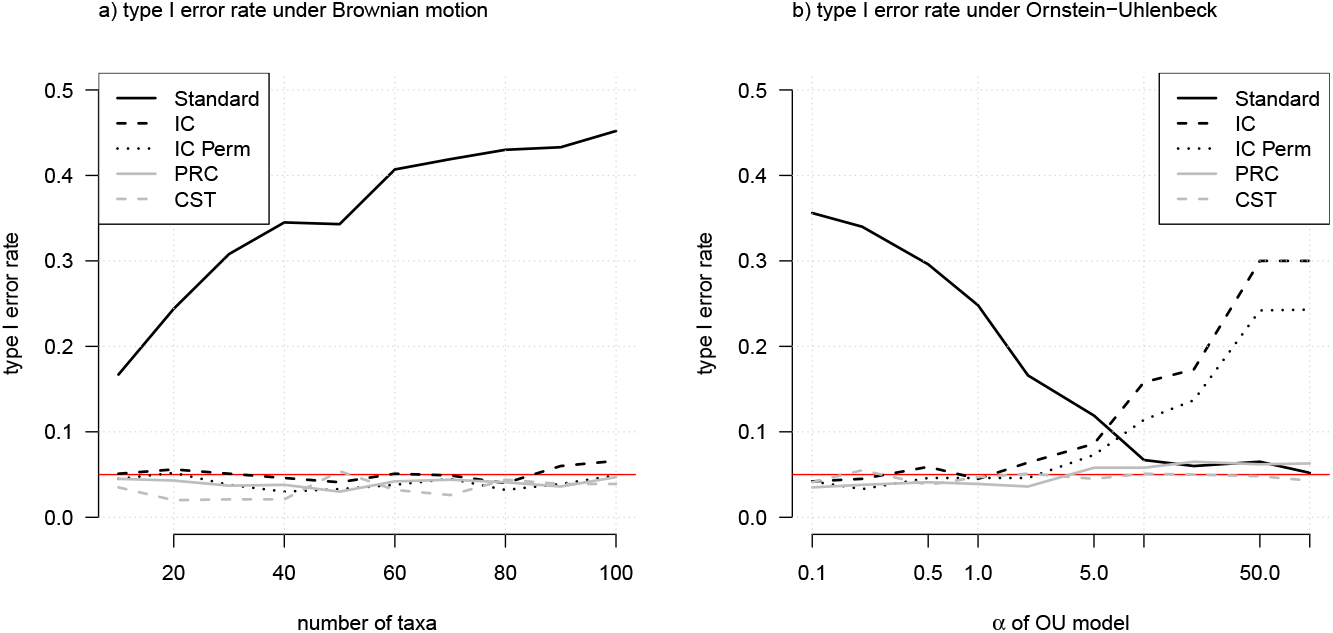
Results of the analysis of type I error. (a) Type I error of both a standard (non-phylogenetic) correlation, and various phylogenetic correlations, when data for *X* and *Y* were simulated under a Brownian motion model, for various total number of taxa, *N*. (b) Type I error for standard and phylogenetic correlation tests when data were simulated under an Ornstein-Uhlenbeck process, for various values of *α*. Legend labels are as follows: “Standard,” non-phylogenetic correlation; “IC,” correlation test based on contrasts; “IC Perm,” same as “IC,” but in which a null distribution of the test statistic was obtained via permutation of the contrasts; “PRC,” phylogenetic rank correlation (of this article); and “CST,” contrasts sign test of Ackerly (2000). See main text for additional details.

We next proceeded to simulate uncorrelated data for *X* and *Y*, but this time under an Ornstein-Uhlenbeck, rather than Browian, process. The Ornstein-Uhlenbeck process is a stochastic process similar to Brownian motion, but in which there exists a tendency to revert towards some central value of the trait (Hansen 1997; Butler and King 2004). (As such Ornstein-Uhlenbeck is sometimes referred to as Brownian motion with a ‘rubber-band,’ in which we can imagine an elastic rubber bands that tends to pull the stochastic process towards its tether-point.) The strength of this tendency is determined by the model parameter *α* (not to be confused with our nominal type I error rate, also denoted as *α*). Higher values of *α* correspond to a greater tendency to revert towards the central value. The parameter *α* can also be understood in terms of “phylogenetic half-life” (Hansen et al. 2008), ln(2)*/α*, which is equal to the average time (under the process) that a trait should require to evolve from its ancestral value halfway to the optimum of the Ornstein-Uhlenbeck process. Conversely, a value of *α* = 0 (and consequently a phylogenetic half-life of ln(2)*/α* = ∞) corresponds to Brownian motion evolutionary change.

Using only the set of 1,000 simulated trees with *N* = 50 from our prior analysis, we simulated data for *X* and *Y* under an Ornstein-Uhlenbeck model with values of *α* = 0.1, 0.2, 0.5, 1, 2, 5, 10, 20, 50, and 100. This resulted in a total of 10 × 1, 000 = 10, 000 additional simulated datasets. We then repeated the same non-phylogenetic and phylogenetic correlation analyses as were described in the prior section.

Results for this test are shown in Figure 2b. In general, we found that type I error of the non-phylogenetic (i.e., standard) correlation was highest for low *α* and decreased with increasing *α* (Figure 2b). This is fairly unsurprising because low *α* corresponds closely to Brownian motion in which a standard correlation is unreliable; while (by contrast) very high *α* tends to reduce the autocorrelation of species data which, in the limit as *α* → ∞, will lose all phylogenetic autocorrelation altogether and thus satisfy the assumptions of standard OLS. Conversely, type I error of all phylogenetic methods was close to the nominal level of *α* = 0.05 for the lowest values of (Ornstein-Uhlenbeck) *α*, but increased for higher *α*. This increase, however, was smallest for the CST and PRC (of this article), and highest for standard independent contrasts and permutation-based contrasts regression (Figure 2b). In particular, type I error at the very *highest* level of *α* was actually slightly below 5% for the CST, 6.3% for the PRC, and around 30% for contrasts regression.

Taken together, these results indicate that our semiparametric contrasts rank correlation method (PRC) has type I error under Brownian motion that is virtually identical to existing parametric and non-parametric phylogenetic evolutionary correlation tests (Figure 2a). Our results also show, however, that the phylogenetic rank correlation test can have much lower type I error than parametric methods when the strict assumption of Brownian motion evolution is violated by our data (Figure 2b).

In addition to measuring type I error, we also wanted to test the power of the PRC method to detect a statistically significant correlation where one exists. To this end, we commenced by simulating data for *X* and *Y* using exactly the same conditions as for Figure 2, described above, but using an evolutionary correlation between the traits of *r* = 0.5. Since the simulations in every other way paralleled our type I error test, the total number of data sets generated for this analysis was 10,000 + 9,000 = 19,000. Power for each simulation condition and method was scored as the fraction of P-values than were equal to or fell below 5%. The results for this analysis are given in Figure 3.

**Figure 3.**
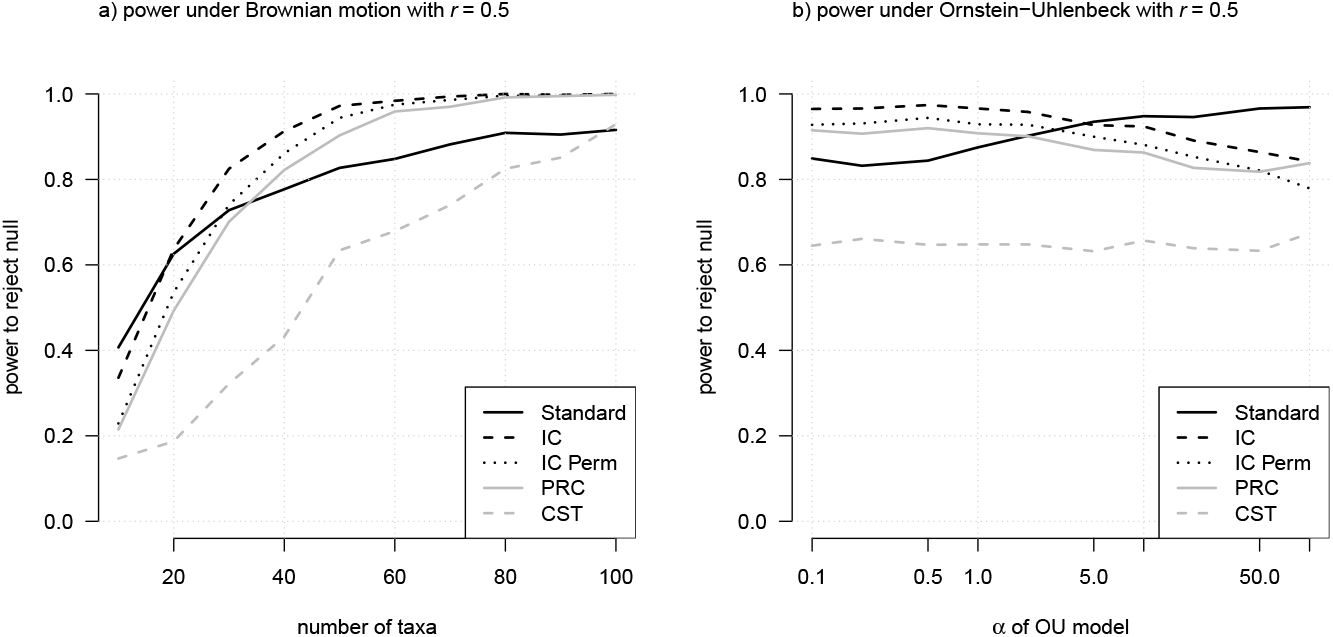
Results of the analysis of power for a true underlying evolution correlation between *X* and *Y* of *r* = 0.5. (a) Power of both a standard (non-phylogenetic) correlation, and various phylogenetic correlations, when data for *X* and *Y* were simulated under a Brownian motion model, for various total number of taxa, *N*. (b) Power of standard and phylogenetic correlation tests when data were simulated under an Ornstein-Uhlenbeck process, for various values of *α*. Legend as in Figure 2.

When *X* and *Y* were simulated under Brownian motion, we found that standard contrasts (IC) and contrasts with a statistical test via permutation (IC Perm) had the highest power; however, these methods were followed closely by our semiparametric PRC (Figure 3a). Ackerly’s (2000) CST method had the lowest power of all methods (Figure 3a). When our correlated data for *X* and *Y* were simulated under Ornstein-Uhlenbeck a similar pattern repeats in which standard contrasts (IC), a permutation test using contrasts (IC Perm), standard (non-phylogenetic) correlation, and our phylogenetic rank correlation (PRC) have the highest power – much higher than the power of the CST by Ackerly (2000; Figure 3b).

Lastly, to explore the influence of effect size on statistical power, we used the same set of 1,000 simulated trees employed for the previous analysis in which the number of taxa *N* = 50. We first simulated correlated Brownian motion evolution on each set of trees while varying the evolutionary correlation coefficient, *r*, between *r* = 0 and 1.0 on the interval 0.1 (*r* = 0, 0.1, 0.2, and so on). We next repeated the same simulation, but using Ornstein-Uhlenbeck with *α* = 10 for simulation. In total, we thus generated a total of 11 × 1,000 × 2 = 22,000 simulated data sets for this analysis.

For each simulation condition, we measured power as the fraction of statistical tests from each simulation condition rejecting the null hypothesis of no correlation (although, of course, this was just a re-measurement of the type I error for *r* = 0). Results for these analyses are shown in Figure 4. In general, when data were simulated under Brownian motion, our PRC test had highly similar power to contrasts regression (Figure 4a). The CRT had much lower power over all simulation conditions (Figure 4a). Standard, non-phylogenetic correlation had high “power” for low values of *r*; however, this actually reflects the much higher type I error of non-phylogenetic methods that we showed in Figure 2a. By contrast, for higher values of *r* the power of non-phylogenetic analyses falls substantially below both the contrasts method and our PRC. Likewise, for data simulated under an Ornstein-Uhlenbeck model with *α* = 10, the PRC method of this article exhibited power across all effect sizes that was substantively higher than the CRT and comparable to both standard (non-phylogenetic) and contrasts-based correlation methods (Figure 4b). Consistent with our analysis of Figure 2b, however, the PRC also showed *lower* “power” (reflecting its lower type I error, and thus a desirable attribute of the method) for the lowest *r* values of *r* = 0 and 0.1 (Figure 4b).

**Figure 4.**
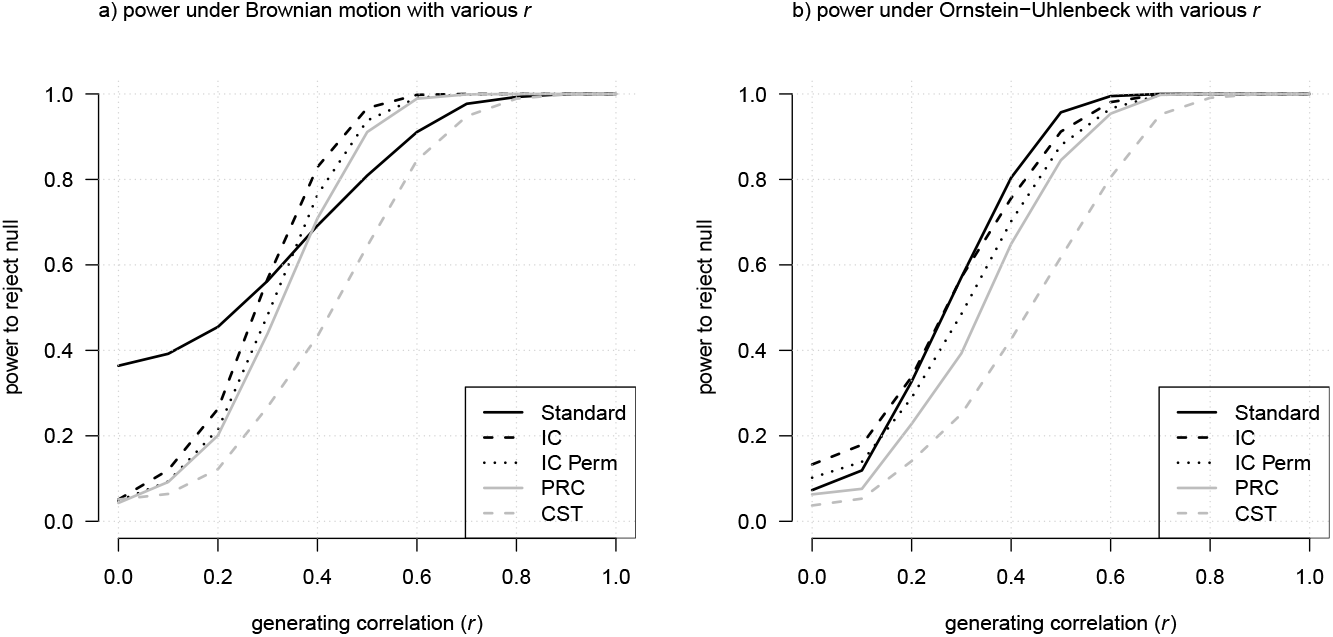
Results of an analysis of power, for various evolutionary correlations between *X* and *Y*. (a) Power of both a standard (non-phylogenetic) correlation, and various phylogenetic correlations, when data for *X* and *Y* were simulated under a Brownian motion model, for evolutionary correlations, *r*. (b) Power of standard and phylogenetic correlation tests when data were simulated under an Ornstein-Uhlenbeck process with *α* = 10, for various values of the evolutionary correlation, *r*. Legend is as in Figures 2 and 3.

## 4 DISCUSSION

Phylogenetic comparative methods have become an indispensable tool of evolutionary research (reviewed in O’Meara 2012; Harmon 2019; Revell and Harmon 2022), and are increasingly employed across a wide variety of different research fields, from community ecology, to cancer biology, to infectious disease epidemiology (e.g., Peterson et al. 2021; Gao et al. 2022; Magalis et al. 2024). Even as they grow in complexity and sophistication, a simple test of the evolutionary correlation of two variables (while accounting for statistical non-independence arising from common descent) remains among the most popular phylogenetic comparative analyses. Typical methods for measuring the evolutionary correlation between traits, however, rely on an assumption of Brownian motion evolution of the variables or their residual error (Felsenstein 1985; Garland et al. 1992).

In this contribution, we introduce a new semiparametric phylogenetic rank correlation (PRC) test that seeks to relax this assumption. We have characterized this method as “semiparametric” because it is not truly distribution-free. Indeed, the method begins via the calculation of standardized independent contrasts (see Figure 1). Standardization is undertaken by dividing each raw contrast by its expected variance, a step that implicitly assumes our traits have evolved under a Brownian process (Felsenstein 1985)! Nonetheless, we assert, and the results of our simulation analyses show, that by converting these contrasts to ranks, the PRC method substantially lessens the effects of violations of this assumption for testing correlations between characters (Figures 2, 3, 4). This is probably because by transforming the standardized contrasts into ranks, our assumption of Brownian evolution effectively changes into the much weaker assumption that longer branches of the tree should contain more evolution of the trait (on average) than do shorter branches. Though this is likely to be true across a much broader swathe of evolutionary processes than is strict Brownian motion evolution, it’s not difficult to envision circumstances in which this assumption (too) is violated. Indeed, we would predict that strong heterogeneity in the rate of evolution (e.g., O’Meara et al. 2006) will lead to elevated type I error of the PRC, just as it would to all of the other phylogenetic methods explored in this article. In a case were substantial heterogeneity of the rate of evolution is thought to exist, we would recommend a method designed to explicitly account for this variation (e.g., Revell and Collar 2009; Caetano and Harmon 2018).

## 5 CONCLUSIONS

Phylogenetic comparative methods are often employed to measure the correlation between continuously-valued species characteristics. Here, we provide a robust semiparametric correlation test that has fewer assumptions than do traditional approaches, but higher power than competing non- or semiparametric methods (e.g., Ackerly 2000). We propose that this method be considered under condition in which the statistical significance of an evolutionary correlation between traits is of interest, but where the satisfaction of model assumptions cannot be assured.

## 6 CODE AND DATA AVAILABILITY

All analyses of this article were undertaken in R (R Core Team 2024), using the contributed R packages *ape* (Paradis and Schliep 2019), *geiger* (Pennell et al. 2014), and *phytools* (Revell 2024). Parallelization of simulations was accomplished using the *foreach* package (Microsoft and Weston 2022). Manuscript drafts of the article were written in Rmarkdown (Xie et al. 2018, 2020; Allaire et al. 2023), and developed with the help of both *bookdown* (Xie 2016, 2023) and the posit Rstudio IDE (RStudio Team 2020). All data and markdown code necessary to exactly rebuild the submitted version of this article (including its analyses and figures) can be found at https://github.com/liamrevell/Revell-and-Harmon.NonParametricPCM.

## 7 ACKNOWLEDGMENTS

We thank J. J. Kolbe, J. B. Losos, D. Schluter, R. E. Glor, B. Banbury, T. Hagey, C. Brock, and H. Alamillo for providing helpful feedback on earlier versions of this project.

